# Bayesian Approach Enabled Objective Comparison of Multiple Human iPSC-derived Cardiomyocytes’ Proarrhythmia Sensitivities

**DOI:** 10.1101/2023.05.14.540739

**Authors:** Tetsuro Wakatsuki, Neil Daily, Sunao Hisada, Kazuto Nunomura, Bangzhong Lin, Ko Zushida, Yayoi Honda, Mahoko Asyama, Kiyoshi Takasuna

## Abstract

The one-size-fits-all approach has been the mainstream in medicine, and the well-defined standards support the development of safe and effective therapies for many years. Advancing technologies, however, enabled precision medicine to treat a targeted patient population (e.g., HER2+ cancer). In safety pharmacology, computational population modeling has been successfully applied in virtual clinical trials to predict drug-induced proarrhythmia risks against a wide range of pseudo cohorts. In the meantime, population modeling in safety pharmacology experiments has been challenging. Here, we used five commercially available human iPSC-derived cardiomyocytes growing in 384-well plates and analyzed the effects of ten potential proarrhythmic compounds with four concentrations on their calcium transients (CaTs). All the cell lines exhibited an expected elongation or shortening of calcium transient duration with various degrees. Depending on compounds inhibiting several ion channels, such as hERG, peak and late sodium and L-type calcium or IKs channels, some of the cell lines exhibited irregular, discontinuous beating that was not predicted by computational simulations. To analyze the shapes of CaTs and irregularities of beat patterns comprehensively, we defined six parameters to characterize compound-induced CaT waveform changes, successfully visualizing the similarities and differences in compound-induced proarrhythmic sensitivities of different cell lines. We applied Bayesian statistics to predict sample populations based on experimental data to overcome the limited number of experimental replicates in high-throughput assays. This process facilitated the principal component analysis to classify compound-induced sensitivities of cell lines objectively. Finally, the association of sensitivities in compound-induced changes between phenotypic parameters and ion channel inhibitions measured using patch clamp recording was analyzed. Successful ranking of compound-induced sensitivity of cell lines was validated by visual inspection of raw data.

**Author Summary:** Cardiac safety is one of the most stringent regulatory risk monitoring during drug development. Regulatory agencies, including the Food and Drug Administration (FDA), require drug developers to conduct thorough preclinical and clinical studies to evaluate drug-induced proarrhythmia risks. The CiPA (Comprehensive in vitro Proarrhythmia Assay) initiative led by the FDA validated the applications of human cardiomyocytes derived from induced pluripotent stem cells in proarrhythmia risk assessments. The assay can be cost effective, and use of high-throughput approaches enables scale-up analysis. However, limited diversity in cell lines to recapitulate heterogeneity of human patients are still lacking in the cardiac safety analysis. We applied various computational tools to maximize capacity for analyzing diverse response of commercially available five cell lines against reference chemical compounds. Limited number of experimental data made it difficult to predict similarities and differences in drug responses among different cell lines. By applying Bayesian approach, our analysis could intuitively grasp the probabilities of expecting different responses from different cell lines.

## Introduction

Cardiac safety is one of the critical concerns among drug developers and regulators to protect against unintended side effects from consumers. At the same time, many drug candidates are not advancing to clinical study due to cardiac safety concerns during preclinical studies [1]. Considering the well-recognized differences in cardiac physiology among species, especially human vs. other animal models, *in vitro* studies using samples representing humans’ genetics and physiology are desired to predict clinical outcomes accurately. Therefore, CiPA (Comprehensive in vitro Proarrhythmia Assay) initiative introduced a new paradigm by which drug-induced cardiac safety is analyzed with *in vitro* assays using human induced pluripotent stem cell-derived cardiomyocytes (hiPSCMs) and *in silico* computational modeling of human adult cardiomyocytes (CMs). One of the advantages of the CiPA approach is its possibility to depict the genetic diversities of human populations through personalizing hiPSCMs and computational models. For example, the analysis of 480 human right atrial tissues isolated from sinus rhythm or atrial fibrillation patients revealed high phenotypic variabilities in action potential (AP) morphology [2], postulating the presence of similar diversities in healthy human CMs. A population *in silico* modeling has already demonstrated its effectiveness in predicting the potential phenotypic diversity of human CMs [3] and proarrhythmic potential [4]. In addition, a recent publication analyzed phenotypes in diverse types of hiPSCMs generated by different laboratories [5], but many cardiac safety studies use a single or few hiPSCM lines, including highly-respected safety consortium studies [6] [7] [8]. While there are potential complications in qualifying hiPSCM lines for cardiac safety evaluations, analyzing diverse cell lines to understand the expected diversity is warranted. However, it is challenging to analyze compound-induced cardiac safety profiles against a diverse panel of hiPSCM lines.

Since 2013, the Consortium for Safety Assessment using Human iPS cells (CSAHi; http://csahi.org/en/), based on the Japan Pharmaceutical Manufacturers Association (JPMA), has been supporting projects in drug safety evaluation using human iPS cell-derived cells, including cardiomyocytes [9]. To address the above-mentioned challenge, here, we used ten known potential proarrhythmic compounds and analyzed their effect on cardiac phenotypes of five hiPSCM lines available commercially. Consequently, the project collected >2,000 time-course traces of calcium transient (CaT) using ten compounds against five cell lines. To analyze the data efficiently and objectively, firstly, we parameterized compound-induced complex changes in CaT traces and clustered. Secondly, Bayesian inference was applied to supplement limited sample replicates that can be used in experiments. Finally, we used canonical correlation analysis to evaluate expected changes in phenotypic parameters against ion channel inhibitions measured using available manual patch clamp data [10]. Prospectively, the combination of Bayesian statistics and experimental analysis, as well as multiparameter analysis, would be useful in high-throughput data analysis with a limited number of samples.

## Methods

### Experiments

#### Cell line and seeding densities

Frozen hiPSCMs vials were thawed from liquid nitrogen in a 37°C water bath for 3 minutes and resuspended in plating media according to each manufacturer’s instructions. hiPSCMs were seeded onto pre-coated (10 ug/mL fibronectin) 384-well low volume plates (Corning, USA) at densities of 4000 cells/well (CDI-iCell^2^ and Miracell) and 8000 cells/well (Axol, Asclestem, and Myoridge). Cells were cultured for 7∼9 days in the culture medium recommended from each manufacturer before compound treatment in a CO2 incubator at 37°C. Each cell line was seeded onto separate 384-well plates.

#### Calcium dye and compound treatment

hiPSC-CMs were stained with a fluorescent calcium dye, EarlyTox^TM^ Cardiotoxicity Kit (Molecular Devices LLC, CA, USA) by incubation for 2 hour in culture medium. Simultaneous fluorescent signals from the entire plate were recorded on an FDSS-µCELL (Hamamatsu Photonics K.K., Shizuoka, Japan) fluorescent plate reader for 5 minutes. A baseline signal was recorded before compound treatment. The cell plate was treated simultaneously with 4 doses of 10 compounds with each dose having 5 replicate samples. After 10 minutes in the above incubator, a compound-treated signal was recorded.

#### Test compounds

Ten compounds and their concentrations used for the analysis is listed in Table 1.

**Table 1.**
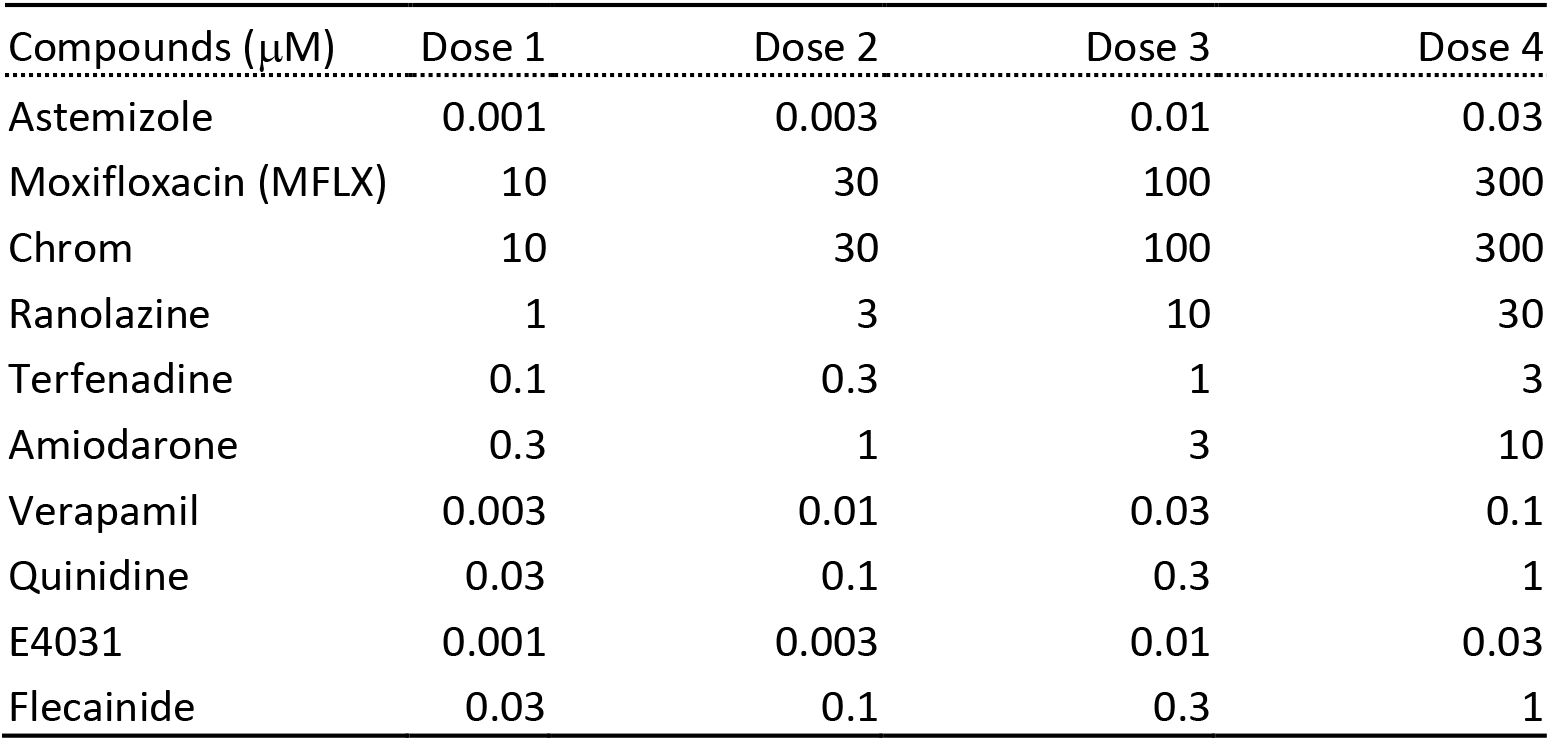
Ten reference compounds and their concentrations used in the assay.

| Compounds (μM) | Dose 1 | Dose 2 | Dose 3 | Dose 4 |
| --- | --- | --- | --- | --- |
| Astemizole | 0.001 | 0.003 | 0.01 | 0.03 |
| Moxifloxacin (MFLX) | 10 | 30 | 100 | 300 |
| Chrom | 10 | 30 | 100 | 300 |
| Ranolazine | 1 | 3 | 10 | 30 |
| Terfenadine | 0.1 | 0.3 | 1 | 3 |
| Amiodarone | 0.3 | 1 | 3 | 10 |
| Verapamil | 0.003 | 0.01 | 0.03 | 0.1 |
| Quinidine | 0.03 | 0.1 | 0.3 | 1 |
| E4031 | 0.001 | 0.003 | 0.01 | 0.03 |
| Flecainide | 0.03 | 0.1 | 0.3 | 1 |

#### Waveform analysis

After automatically finding all the CaT cycles using iVSurfer (InvivoSciences, Inc, Madison, WI, USA). The six parameters including peak-width duration 80 (P80), peak-width duration 80/20 (8/2), cycle-length (CL), amplitude (AMP), number of peaks in the recorded timeframe (PN), and standard deviation of cycle lengths of all peaks (CSD) were computed. Each parameter was calculated as a percent change from the baseline (before compound addition). The 8/2 was calculated dividing P80 by P20. Because three times P80 over P20 often signified an early after depolarization (EAD), we used a sigmoidal function, (exp(-9(x-3)+1)^-1^. CL was equal to the duration between two peaks. PN was the number of CaT cycles found within the recording (185 seconds). Standard deviations of CL values in each recording, CSD, represented variance of CL, indicating instable CaT wave trains. The log-modulus transformation: *L*(*x*) = *sign*(*x*)log (|*x*| + 1) [11] was applied to visualize a wide range of parameter values. By plotting dose-dependent relative changes of six parameters in radial plots, the trends in compound-induced changes to parameters were visualized to compare cell lines’ differences and similarity of compound-induced responses.

#### Bayesian and PCA analysis

Bayesian approach was applied to estimate sample distributions of each compound-induced data set (n=5). Sample number was increased to 1,000 with Bayesian bootstrapping. The Bayesian bootstrap using Dirichlet distribution produced a more uniform gaussian distribution than classic bootstrapping [12].

Principal component analysis and hierarchical clustering analyzed compound-induced changes in calcium transient waveforms defined by 24 features; six parameters (P80, 80/20, CL, AMP, PN, and CSD) in four doses. To estimate the statistical variance in each parameter, Bayesian bootstrapping simulated the posterior distribution of 1000 samples using experimental data as a priori.

#### Canonical correlation analysis

Canonical correlation analysis (CCA) of the dataset was performed in MATLAB using the canonical correlation function: canoncorr. CCA was performed between the phenotypic dataset and the ion channel inhibition dataset for each compound of each cell line.

## Results

### Cell lines with ten drugs

Average profiles of CaTs of five commercially available hiPSCM lines, beating spontaneously, were analyzed by calculating their ensemble averages of all the cardiac cycles (Figure 1A-E). The wavelet transform algorithm automatically detected all the CaT cycles in various beating rates of five hiPSCM lines. While their spontaneous rates of CaT were consistent within each cell line, further analysis revealed their significant differences in spontaneous beat rates (i.e., the inverse of cycle lengths) and CaT profiles among different cell lines. For example, Myoridge cells had the shortest CaT duration with a quick rise/decline and were beating faster than others (Figure 1E). By analyzing the effects of compounds on CaTs, four concentrations of ten drugs with known potential proarrhythmic risks (Table 1) were applied to the hiPSCMs growing in 384-well plates using protocols and media provided by each vendor. Each well received a single dose of a drug, and CaT traces before and after drug treatments were compared. As we and others [1, 13] utilized durations of CaTs as a key metric to gauge drug-induced proarrhythmia sensitivity of hiPSCMs, dose-dependent changes in CaT duration average of all cell lines were plotted (Figure S1). Most of the hERG inhibitors increased CaT duration significantly, as expected. However, by inspecting individual CaT traces, we found that some cell lines stopped beating or seldom but consistently exhibited irregular beats at some dose levels (Figure 1G). Therefore, simply analyzing only CaT duration or beat rates could miss those proarrhythmic events. Consequently, the diligent observations of raw data traces of many cell lines pointed to the importance of multidimensional data analysis.

**Figure 1.**
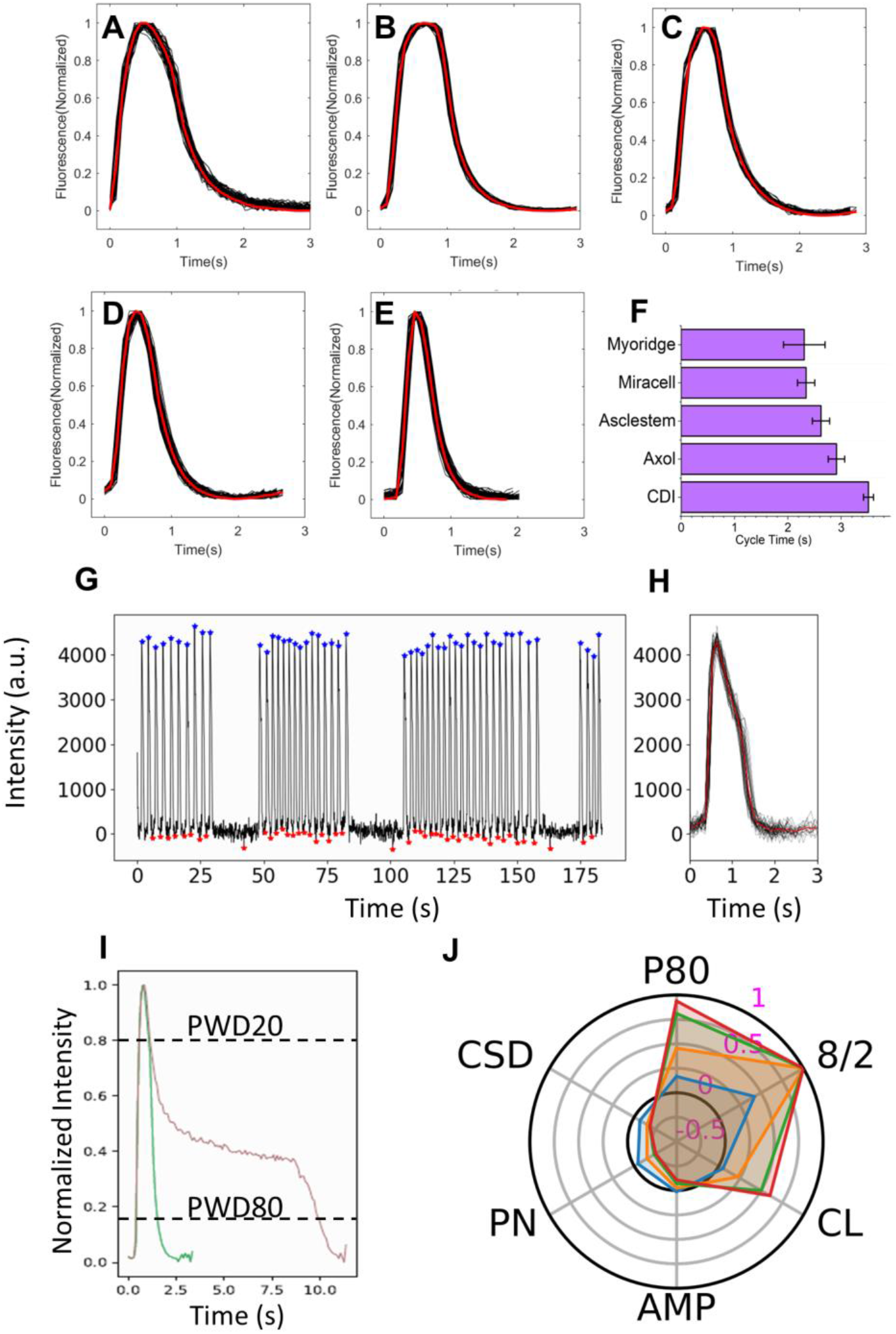
Calcium transients of five hiPSCM cell lines and multi-parameter phenotyping. Ensemble average (red trace) of calcium transients (CaTs, black trace) for (A) CDI, (B) Axol, (C) Asclestem, (D) Miracell, and (E) Myoridge cardiomyocytes and their average cycle times (F) were calculated. An example CaT of Myoridge cardiomyocytes after 10 nM E4031 treatment (G) its ensemble average (H) had an irregular beats. The software automatically detected peaks (blue dots) and valleys (red dots). (I) CDI’ CaTs of pre (green) and post (red) E4031 (30 nM) treatment were plotted with PWD80 and PWD20 lines. (J) Radial plot of parameters calculated from CDI cardiomyocytes treated with E4031. Parameters are PWD80 (P80), PWD80/PWD20 (8/2), cycle length (CL), amplitude (AMP), peak number (PN), and standard deviation of the cycle length (CSD). Increasing concentrations of compounds were color-coded polygons as the following: dose1 (blue), dose2 (orange), dose3 (green) and dose4 (red).

### Six-parameter analysis

Multiple parameters were used to define differences in profiles of action potential and CaT previously [14]. We applied a similar approach to parameterized profiles of CaTs and their irregular beating to classify different trace patterns. For analyzing >2,000 traces of each with ∼10,000 time-points, we utilized a data analysis software specialized in periodic data (see method), which automatically detected regular and irregular cardiac beats (Figure 1G) and calculated their ensemble averages (Figure 1H). In addition to the CaT duration (PWD80), five more parameters classified different profiles of CaTs and beat patterns. Among them, ratios of PWD80 over PWD20, namely triangulation indicator [15], were useful for detecting traces exhibiting EAD when it got significantly high (Figure 1I). The number of peaks in an observation period (180 sec) usually is inversely proportional to the cycle length. However, this reciprocal relation does not hold when beats become irregular. For example, some samples showed reduced beat rates without prolonging the CaT durations (Figure S2); samples with irregular beats had increased standard deviation of cycle lengths (Figure S2). Finally, the changes in the six-dimensional parameters of before and after drug treatment (after drug/before drug-1) were plotted radially to visualize compound-induced response patterns (Figure 1J).

### Six-parameter Phenotypic Responses of E4031

One of the major classes of proarrhythmia compounds is an hERG inhibitor. An experimental class III anti-arrhythmia drug, E4031, inhibiting the hERG channel with high specificity, was utilized to visualize drug-induced phenotypic response patterns using the multiparameter approach. E4031 significantly prolonged the duration of CaT and induced EADs in all cell lines with various ranges (2-5 fold increase, Figure 2 A-F). Miracell was the most sensitive to E4031 and stopped beating at its highest concentration (Figure 2E). Asclestem and Axol also elongated their CaT duration, but the PWD80/20 ratios did not increase as much as CDI, which was confirmed by their ensemble average traces (Figure 2A, B, C). We also noticed that E4031 not only elongated Myoride’s CaT duration but also increased cycle length SD (Figure 2G). Among all, Myoridge was the least sensitive to E4031; PWD80 did not change much until adding the highest dose of E4031 (Figure 2F, K). Nonetheless, the radial plots of multiple parameters visually captured the different response patterns of five cell lines. Importantly, the response patterns of multiparameter radial plots correctly represented the drug-induced changes in beat patterns, providing an intuitive sense of how five cell lines responded to E4031.

**Figure 2.**
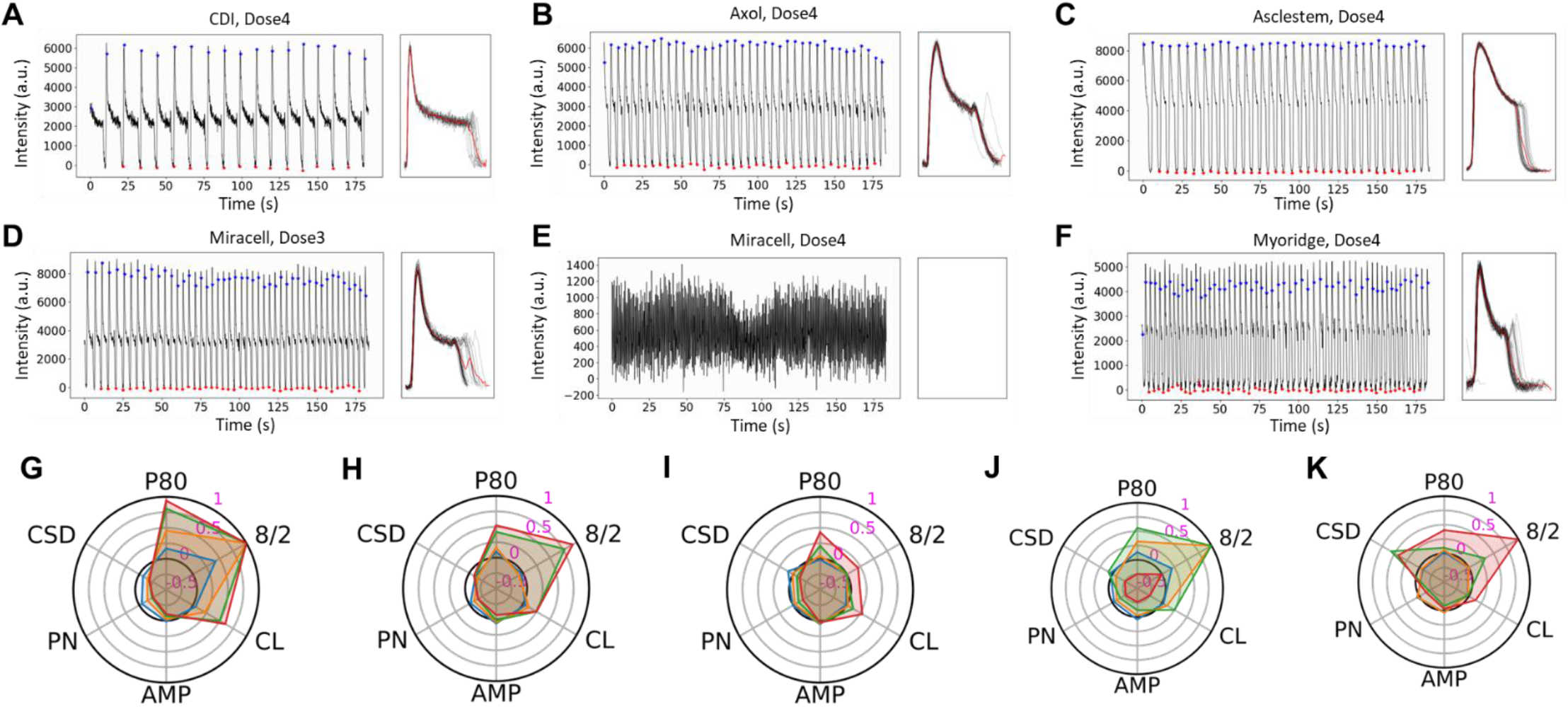
E4031 treatment of hiPSCM cell lines. The CaT analysis, their ensemble average, and comparison to post (red line) and pre (green line) E4031 treatment (30 nM) on cardiomyocytes from (A) CDI, (B) Asclestem, (C) Axol, (D, E) Miracell (10 and 30 nM, respectively) and (F) Myoridge. Analyzed parameters were plated in radial plots for (G) CDI, (H) Asclestem, (I) Axol, (J) Miracell, and (K) Myoridge hiPSCMs.

### Phenotypic Response Patterns between hERG channel inhibition

To further evaluate the benefits of the six-parameter analysis, the effects of other compounds, Astemizole, Flecainide, and Quinidine that chiefly inhibit hERG channel were analyzed; Flecainide and Quinidine marginally inhibit delayed rectifier and transient outward potassium current, IKs and Ito respectively (Figure S14). Their radial plots (Figure 3A-C) revealed similar response patterns – namely, these compounds significantly prolonged CaT duration of CDI and Miracell but not Asclestem (except for Flecainide), Axol, and Myoridge. The other hERG inhibitor, Ranolazine, also inhibiting the late sodium channel, reduced effects on elongating CaTs in CDI and Miracell (Figure 3D), which is in line with its known balancing effects of elongation and shortening of CaT by the dual channel inhibition. Interestingly the highest dose of Ranolazine reduced the amplitude of all cell lines (Figure 3D, S5), which was expected by the mechanism of action as an anti-angina drug to reduce calcium concentration via inhibiting the late sodium current [16]. Additionally, when IKs was inhibited as much as the hERG channel with Moxifloxacin, a fluoroquinolone antibiotic with a slight inhibition of the late sodium current known to prolong QT, [17] (Figure 3E), CDI and Miracell became almost insensitive, but other lines showed prolonged CaT duration or stopped their beating.

**Figure 3.**
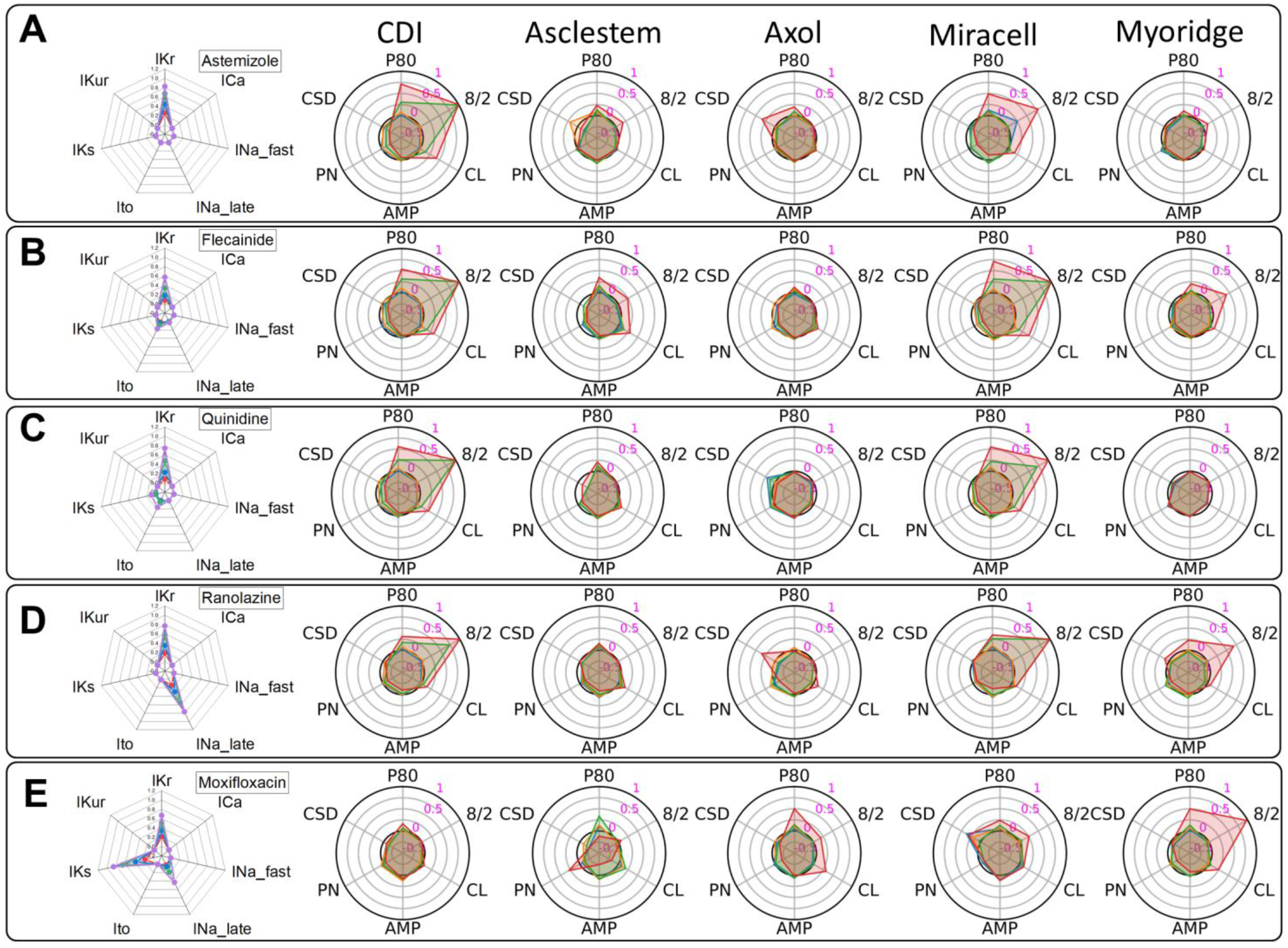
Compounds inhibit IKr of hiPSCM cell lines. Astemizole (A), flecainide (B), quinidine (C), ranolazine (D), and moxifloxacin (E) inhibit rapid delayed rectifier K+ channel (*I*^Kr^) and other ion channels, including L-type Ca2+ channel (*I*_Ca_), fast voltage-gated Na+ channel (*I*_Na fast_), late voltage-gated Na+ channel (*I*_Na late_), transient outward K+ current (*I*_to_), delayed rectifier potassium channel (*I*_Ks_), and ultra-rapid outward current (*I*_Kur_) at various degrees. Radial plots (left in each panel) display published patch-clamp recording of dose-dependent channel inhibition (1.0 = 100% inhibition) with a color code: dose1 (red), dose2 (blue), dose3 (green), dose4 (purple). Radial plots (right) represent the effect of each compound on six parameters of CDI, Asclestem, Axol, Miracell, and Myoridge hiPSCMs.

### Ca channel inhibition and multi-channel inhibition

The radial plot visualization revealed cell line-specific sensitivity features unique to the L-type calcium channel inhibitors with or without inhibiting other channels. We used Verapamil & Terfenadine inhibiting L-type calcium (CaV1) and hERG channels and Amiodarone inhibiting fast and late sodium channels and Ito current in addition to hERG and CaV1 channels. All these drugs inhibited the hERG channel to a different extent; however, almost no prolongation of CaT duration was recorded (Figure 4A-C). Asclestem and Myoridge lines first increased cycle lengths and their SDs (i.e., irregular slow beats) and then stopped beating at higher concentrations. Finally, when Chrom293B (Chrom) inhibiting IKs, Ito, and IKur (ultra-rapid outward current) with marginal IKr but without affecting sodium and calcium channels was applied, all cell lines showed slight prolongation of CaT duration and stopped beating at the highest concentration in some cell lines (Figure 4D).

**Figure 4.**
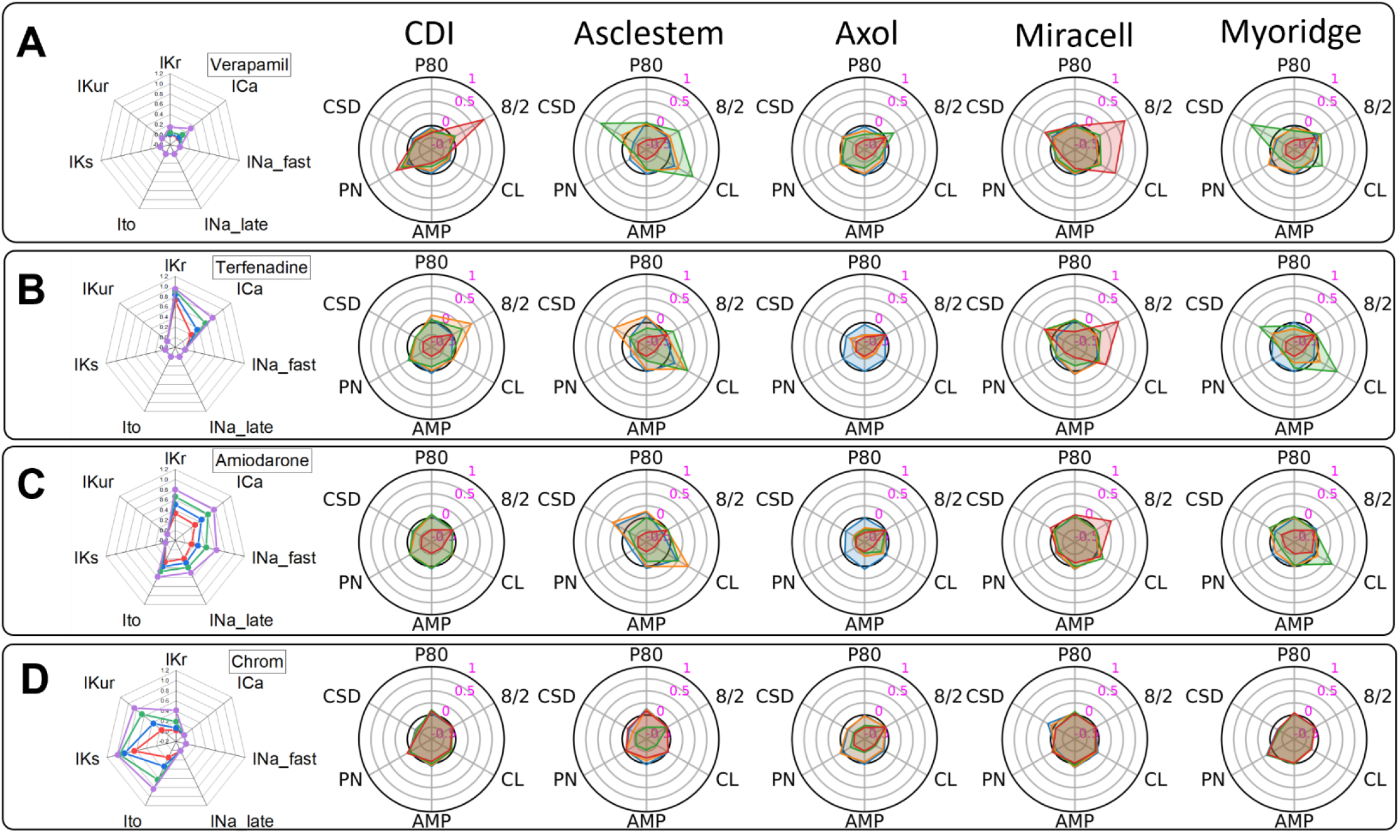
Compounds inhibit L-type Calcium and other channels of hiPSCM cell lines. Verapamil (A), terfenadine (B), and amiodarone (C) inhibit L-type Ca2+ channel (*I*_Ca_) and other channels at various degrees. Chrom (D) inhibits various potassium channels. Radial plots (left in each panel) display published patch-clamp recording of dose-dependent channel inhibition (1.0 = 100% inhibition) with a color code: dose1 (red), dose2 (blue), dose3 (green), dose4 (purple). Radial plots (right) represent the effect of each compound on six parameters of CDI, Asclestem, Axol, Miracell, and Myoridge hiPSCMs.

### Classification of cell lines

The six-parameter polar plots visualized trends in similarities and differences of five cell lines responding to different drugs. However, trend recognition can be subjective; therefore, we applied hierarchal clustering and principal component analysis (PCA) in five replicates of phenotypic data that are described by 24 parameters (four doses of six phenotypes). After the analysis and dimensional reduction, two-dimensional data was visualized to compare cell lines’ compound sensitivity and response reproducibility (Figure S3). In many cell lines, the five replicates of each drug response formed groups, but some did not form well-identifiable clusters. A limited number of experimental replicates is challenging to observe clear cluster formations. Then, we applied the Bayesian bootstrapping [18] approach to simulate the posterior distribution of the phenotypic parameters. After this process, the clusters of ten drug responses were clearly visible in different cell lines (Figure 5A-J). The histograms of data in the first and second principal components showed the distribution of data points as well as their overlaps and separations. A tri-axial ellipsoid with semi-axes of three times the standard deviation in the first, second, and third principal components can represent tightness of data distribution (Figure S4); the smaller the volume, the better assay reproducibility. The sum of ellipsoid volumes indicates CDI’s significantly high drug response reproducibility compared with the other cell lines (Figure 5K). In addition, among all the clusters, CDI samples were well-separated from those of DMSO control, indicating its high sensitivity to our selected ten potential proarrhythmic compounds. The clusters of compounds blocking IKr (E4031, Astemizole, Quinidine, Flecainide, and Ranolazine) were well separated from DMSO control in CDI (Figure 5A). However, our visual observation suggests that the other cell lines’ same drug data clusters were near or indistinguishable from the DMSO cluster. Meanwhile, the cluster of Chrom is near the DMSO in CDI, but that of other cell lines, especially Asclestem and Myoridge, was well-separated from the DMSO cluster.

**Figure 5.**
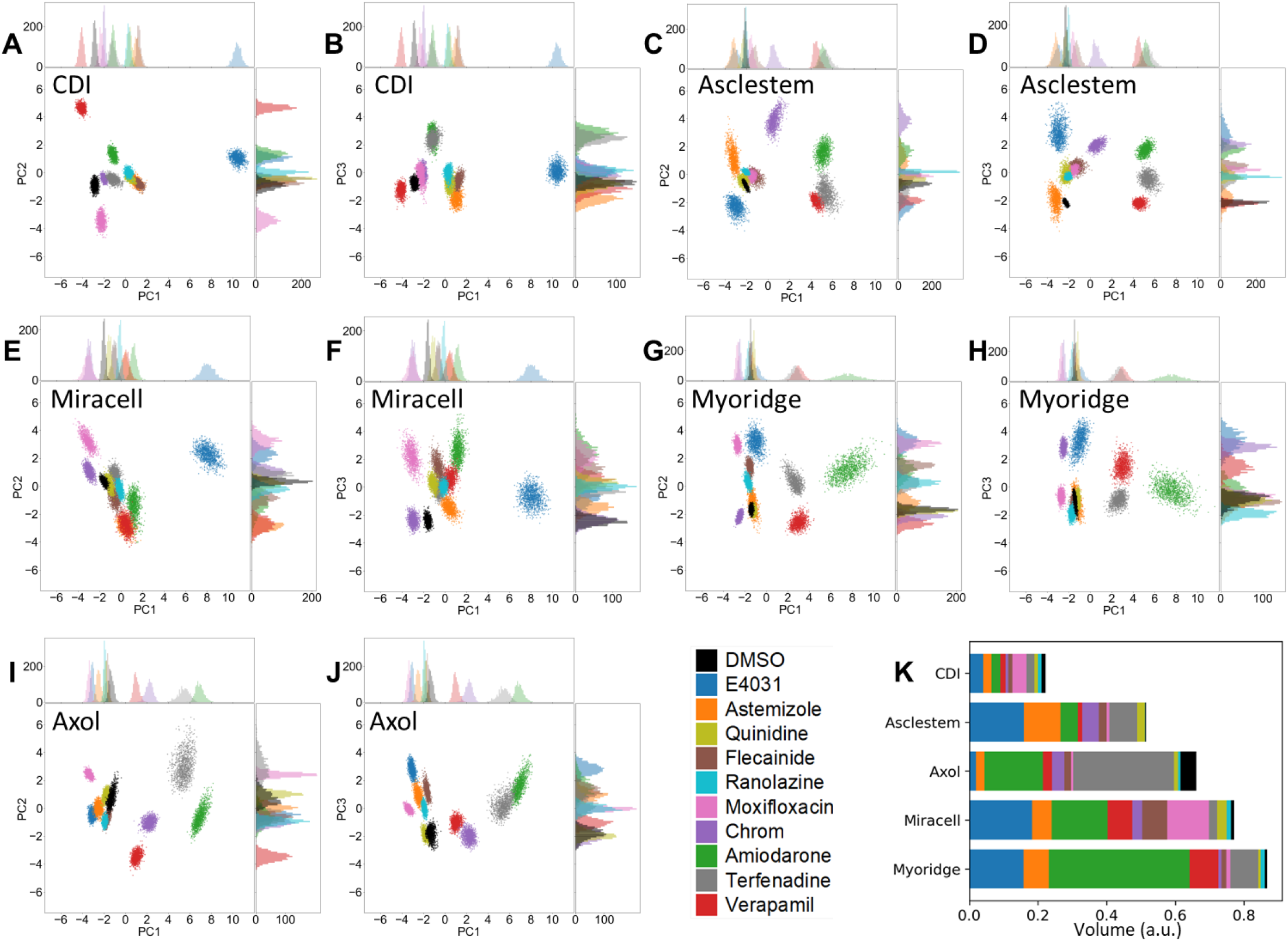
Visualization of compound response data clusters in five hiPSCM lines. After performing principal component analysis of 24 phenotypic parameters, distributions of compound-induced changes in CDI (A, B), Asclestem (C, D), Miracell (E, F) Myoridge (G, H), and Axol (I, J) were plotted. Data were plotted two dimensionally with the first (PC1) and second (PC2) principal components (A, C, E, G and I) and the first (PC1) and third (PC3) principal components (B, D, F, H and J). (K) Volumes of all clusters, calculated by multiplying standard deviations of the first 3 principal components, were stacked to compare sizes of data variance.

### Association between ion channel inhibition and phenotypic response

Drug-induced phenotypic responses in the six parameters were expected to correlate with the inhibition of multiple ion channels inhibited by each compound. CiPA’s study revealed considerable variability in the blocking potency of 12 blinded drugs measured using various automated patch-clamp (APC) at multi-sites [19] and resulted in high uncertainty in the Hill coefficients [20]. Based on the expected large variance in ion channel inhibition data, we simulated the posterior distribution of ion channel blocking using Bayesian bootstrap using IC50 values and Hill coefficients of reference compounds obtained using the manual whole cell patch clamp technique [10]. By assuming a linear correlation between sets of Bayesian-bootstrapped phenotypic parameters and ion channel inhibition, the canonical correlation analysis (CCA) was used to analyze their degree of association (Figure 6A). All the calculated correlation coefficients are near 1.0, indicating strong associations (Figure 6B). However, Ranolazine and Terfenadine had a lower association in all cell lines than other compounds, which indicates that the expected sensitivity of observing phenotypic changes based on the ion channel inhibition data was diminished. In addition, Axol and Myoridge exhibited lower CCA associations for Quinidine compared to other cell lines, which was confirmed by visual inspection of CaT profiles (Figure S6).

**Figure 6.**
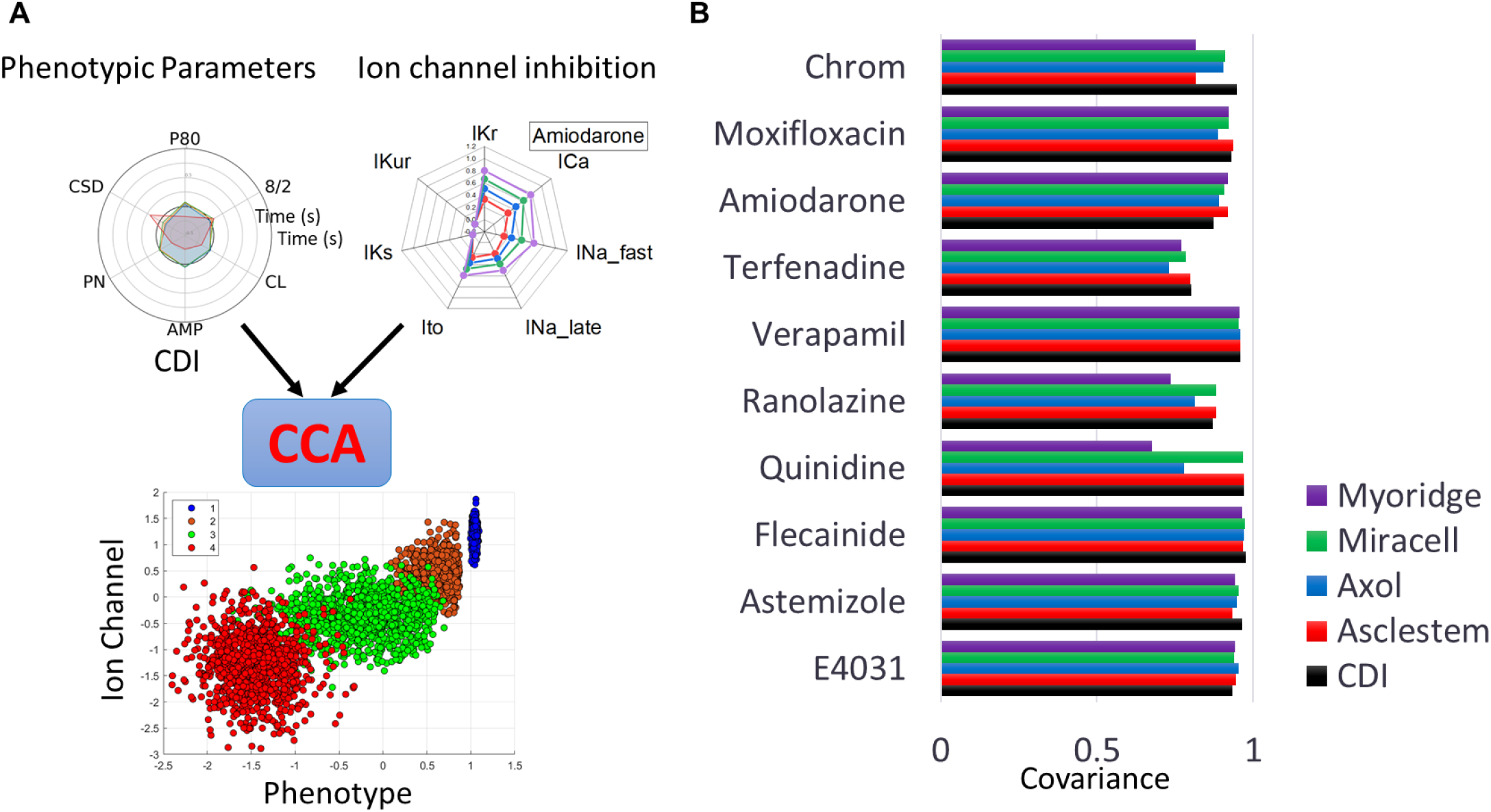
Canonical correlation analysis (CCA) of phenotypic parameters and ion channel inhibition. (A) Diagram showing CCA between phenotypic parameters and ion channel inhibition data. The CCA scatter plot shows the correlation between the first pair of canonical covariates: CCX_1_ and CCY_1_. CCX_1_ is the first canonical covariate for the phenotypic data and CCY_1_ is the first canonical covariate for the ion channel inhibition data. Each dose is plotted by a separate color: dose1 (blue), dose2 (orange), dose3 (green), dose4 (red). The correlation ecoefficiencies between the first pair of canonical covariates for each drug and each cell line were compared.

## Discussion

Six-parameter analysis of calcium traces enabled the classification of five hiPSCM lines treated with ten potential proarrhythmia drugs based on their phenotypic drug responses. The radial plotting of multiparameter data was useful in visually recognizing similarities and differences. However, some of the quantitative data analysis tools enable confirming the visual impressions objectively. In addition, quantitative tools are useful for analyzing large data sets where investigators cannot inspect all the raw data or radial plots. For instance, the canonical correlation analysis can objectively detect the non-responsiveness of Axol and Myoridge cells to Quinidine.

Many lines of CMs have been differentiated from various iPSC lines using different methods [21] [22] by numerous academic laboratories and commercial institutions. Improving the efficacy of CM differentiation and purification reduced the difficulties of their batch production from individualized human iPSCs. Not only for the safety studies but preclinical applications of personalized iPSC-derived samples are advancing the development of personalized medicine (i.e., precision medicine) [23]. In the meantime, it is highly productive to rely upon a well-characterized specific cell line to generate standard data and calibrate diverse test approaches. However, giving too much weight to single cell line data may miss cardiac safety issues caused by the diversity of human genetics. For example, ion channel expressions in CMs are different depending on their locations and even their physiological state [24]. In fact, the pharmaceutical cardiac safety community has been investigating the diversification strategy in proarrhythmia sensitivity analysis, which is successful *in silico* modeling [25]. The diversification in non-virtual assays, however, is expensive and can be time-consuming in analyzing enormous amounts of data. Thus, we introduced the automating of a multiparameter phenotype capturing tool and applied data science tools, including Bayesian bootstrapping, to classify the drug-induced changes to overcome these difficulties. Bayesian bootstrapping is becoming an increasing popular approach in estimating uncertainty of statistical models such as assessing phylogenetic confidence [26] and modeling COVID19 data [27]. In this study, we used it to classify various hiPSCM lines with estimated uncertainty. The dendrograms of the phenotypic response of Axol and Myoridge cells to inhibitors of hERG channel, E4031, Astemizole, Flecainide, and Quinidine were similar and clustered in a group (Figure S7 – S10). Those cell lines also formed a cluster when the late sodium channel, together with the hERG channel, was inhibited by Ranolazine (Figure S11). In general, Axol and Myoridge cells were not sensitive to those inhibitions to change their CaT profiles. Moxifloxacin inhibited late INa, IKs, and IKr, but Axol and Myoridge formed a cluster but were more responsive than the other cell lines (Figure S12, 3E). So, Axol and Myoridge could be a valuable tool for analyzing the mechanism by which Moxifloxacin induced arrhythmia [28]. CDI and Miracell were the most sensitive to hERG inhibitors. A specific L-type calcium channel inhibitor, Verapamil, lowered CaT amplitude of CDI and Axol and eventually stopped their beating but increased cycle length without prolonging CaT duration in Asclestem, Miracell, and Myoridge. A similar trend was seen in Terfenadine and Amiodarone which inhibit IKr and sodium channels in addition to L-type calcium channels. It is interesting to note that Verapamil and Amiodarone reduce heart rate in patients with atrial fibrillation [29, 30] [31], concurring slowing down beat-rate of Asclestem, Miracell, and Myoridge cells

Terfenadine, was removed from the market due to its proarrhythmic risk for long QT-related Torsades de Pointes (TdPs) [32]. The study data using isolated rabbit heart and human atrial myocyte showed that Terfenadine induced non-TdP like ventricular tachycardia/ventricular fibrillation due to a slowed conduction via blockade of *I*Na [33]. The six-parameter phenotypic analysis showed marginal prolongation of CaT duration in CDI cells, but Terfenadine slowed beating of Asclestem, Miracell, and Myoridge and eventually stopped their beating (Figure S13) Axol stopped beating after the second dose. Finally, our analysis revealed that Amiodarone, a class III drug, also slowed the beating of Asclestem with a moderate prolongation of CaT duration. The drug is known to slow heart rate and atrioventricular nodal conduction and prolongs refractoriness [34].

## Limitations

The project focused on determining cell line differences and ranking their sensitivities to different compounds. Therefore, experiments used lower concentration ranges of test drugs to avoid recording without signals (stop beating). However, for analyzing the adequacy of cell lines in safety standard assays, the analysis should use CiPA compounds with their standard concentration range. The data reproducibility of CDI was significantly high among five cell lines. Compared with other cell lines, experimenters are familiar with handling CDI cells. Hence, if protocol and seeding density of other cell lines were optimized, the results could have been different. We were tempted to analyze data further using gene expression analysis of each cell line. However, we could not draw a reasonable conclusion at this time.

## Conclusion

Six-parameter radial plots of CaTs and their quantitative analysis using Bayesian bootstrapping effectively analyzed similarities and differences in five cell lines intuitively and objectively. Multi-parameter approach enabled recognizing various c0mpound-indcued phenotypic changes, including those slowed beat rates without prolonged CaT durations. The canonical correlation analysis ranked expected responsiveness of each iPSCM line to a compound based on its ion channel inhibitions measured by patch clamp recordings. The application of Bayesian approach in high-throughput data analysis where the limited sample size can be challenging in estimating uncertainties and assessing significance of hits. In addition, the application of data science tools facilitated the improvement of the productivity of high throughput data analysis by providing a quantitative framework in decision making. The advances in computational power have made it more feasible for applying computationally intensive approaches, aspiring to support future individualized cardiac safety analysis.

---

**Figure S1.**
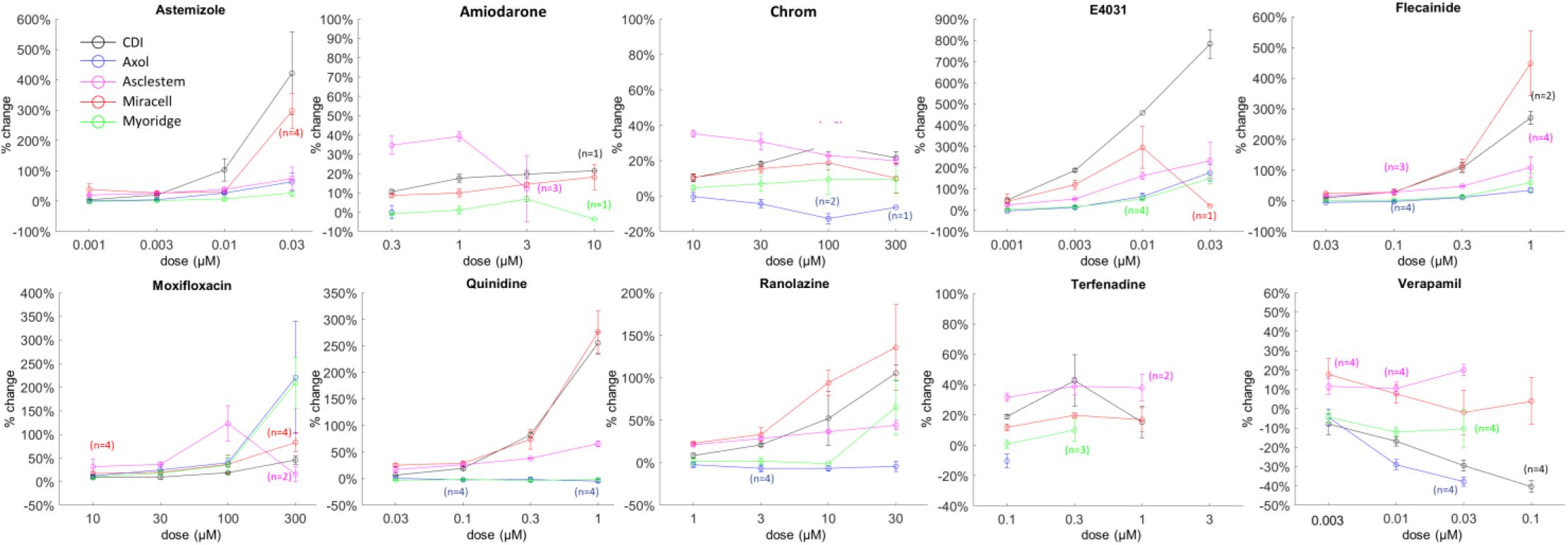
Pulse-width duration 80 (PWD80) analysis of hiPSCM cell lines. Pulse-width duration 80 (PWD80) was calculated by measuring the duration of the calcium transient at 80% from the maximum value. Values are calculated as percent change (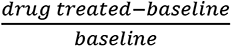) of PWD80. Astemizole increased PWD80 (>200%) in CDI and Miracell. E4031 increased PWD80 (>200%) in CDI, Miracell (< 0.03 µM) and (>100%) in Asclestem, Myoridge and Axol. Flecainide increased PWD80 (>200%) in Miracell and CDI, and (<100%) in Asclestem, Myoridge, and Axol. Moxifloxacin increased PWD80 (>100%) in Asclestem, Axol and Myoridge and (<100%) in Miracell and CDI. Quinidine increased PWD80 (>200%) in Miracell and CDI, and (<100%) in Asclestem, Axol and Myoridge. Ranolazine increased PWD80 (>50%) in CDI, Miracell and Myoridge and (<50%) in Asclestem and Axol. Verapamil decreased PWD80 in CDI, Myoridge and Axol.

**Figure S2.**
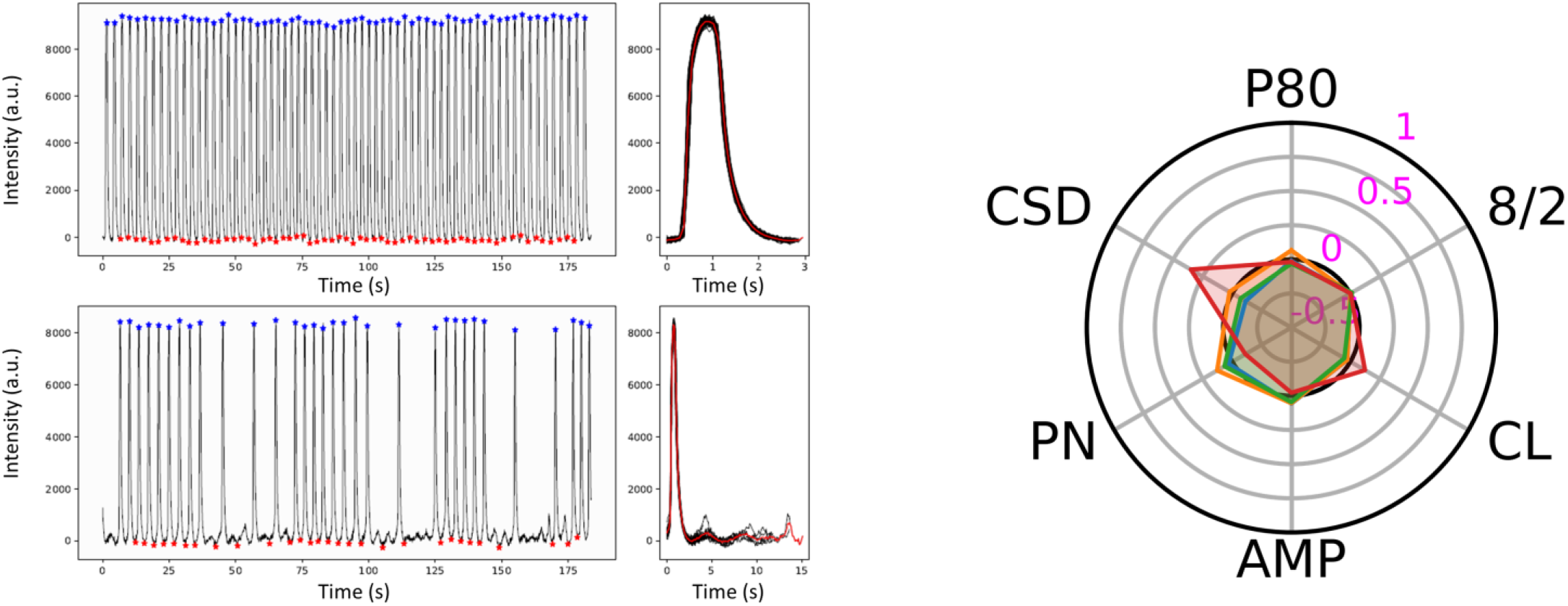
Ranolazine (30 μM) on Axol induced irregular beats without prolonging CaT duration (P80). The radioplt showed no changes in P80 but increased cycle length (CL) and their standard deviation CSD).

**Figure S3.**
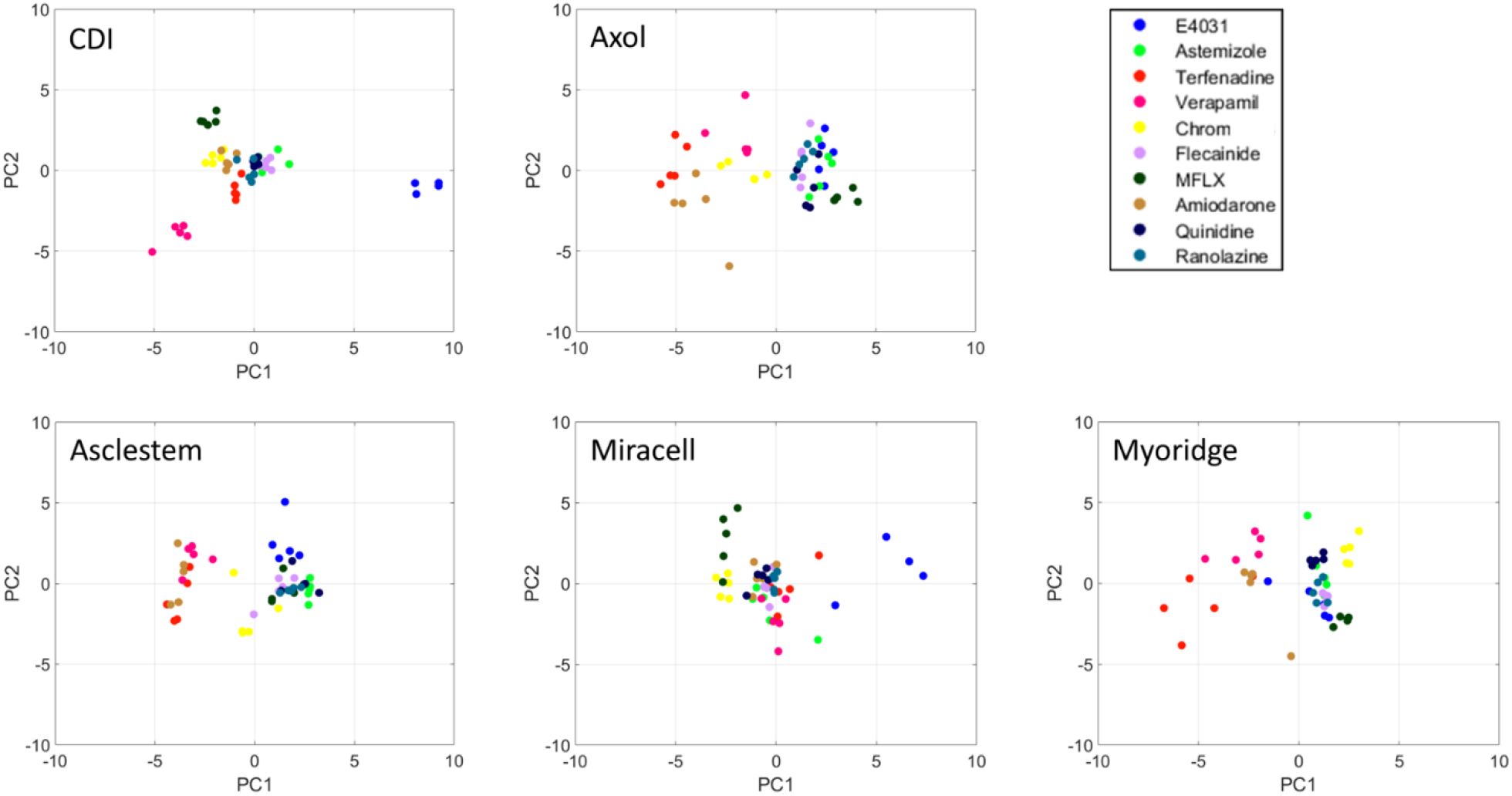
2D plot of raw data after PCA. Five sample data after PCA were plotted on the 1st and 2nd component plane. Data treated by the same compounds seem to form culsters but their distribution were not clearly visible.

**Figure S4.**
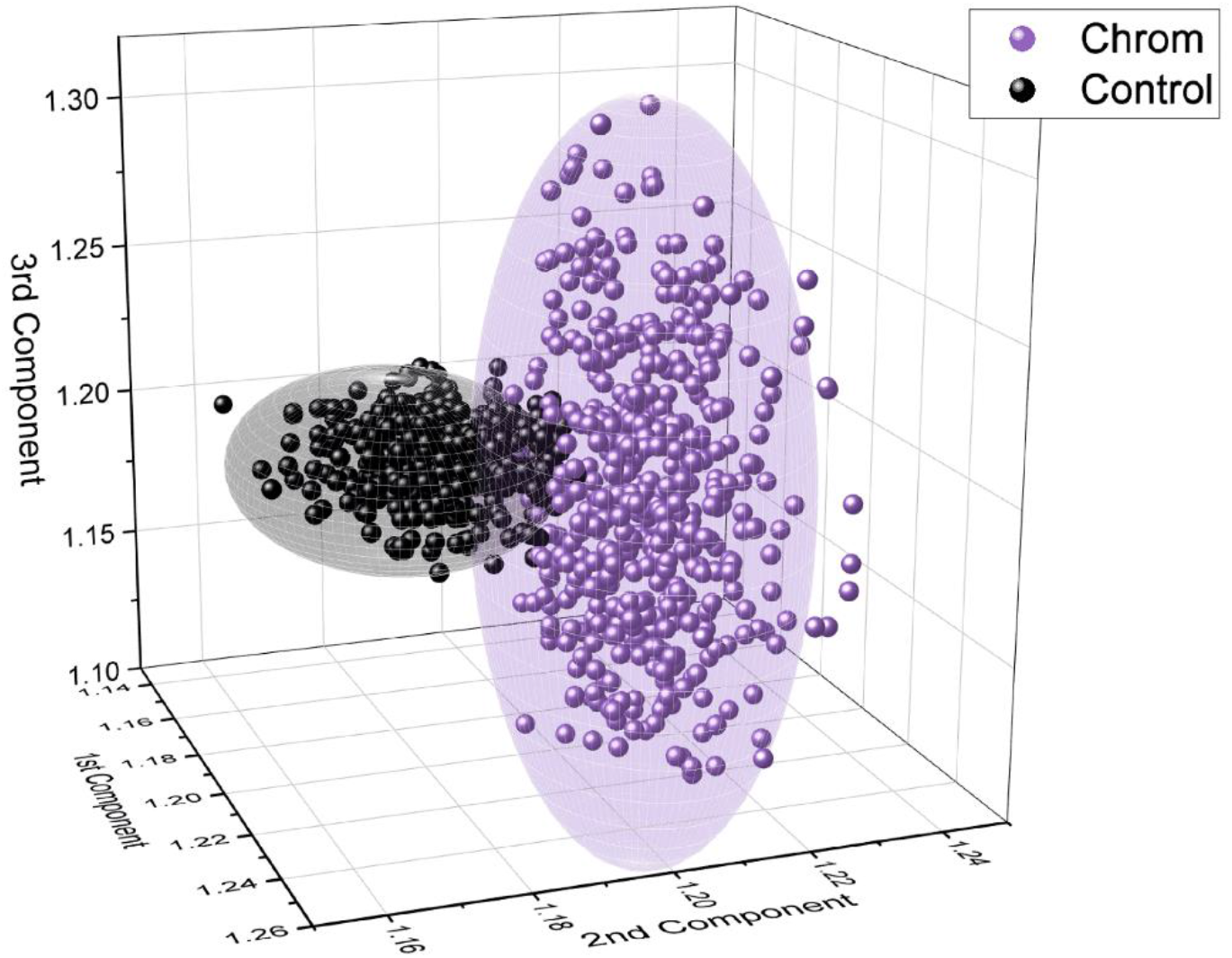
PCA analysis data in three-dimensional space. The 1st, 2nd, and 3rd Component represents each axis of the first three principal components. After Bayesian bootstrapping, DMSO control (1,000 black spheres) and chromanol 293B (Chrom, 1,000 purple spheres) data formed clusters in the 3D space. Standard deviation (SD) of each component axis of control and compound-treated data defined ellipsoids for control and chromanol 293B, enclosing data points. Sizes of ellipsoids represented data distribution with different SD sizes.

**Figure S5.**
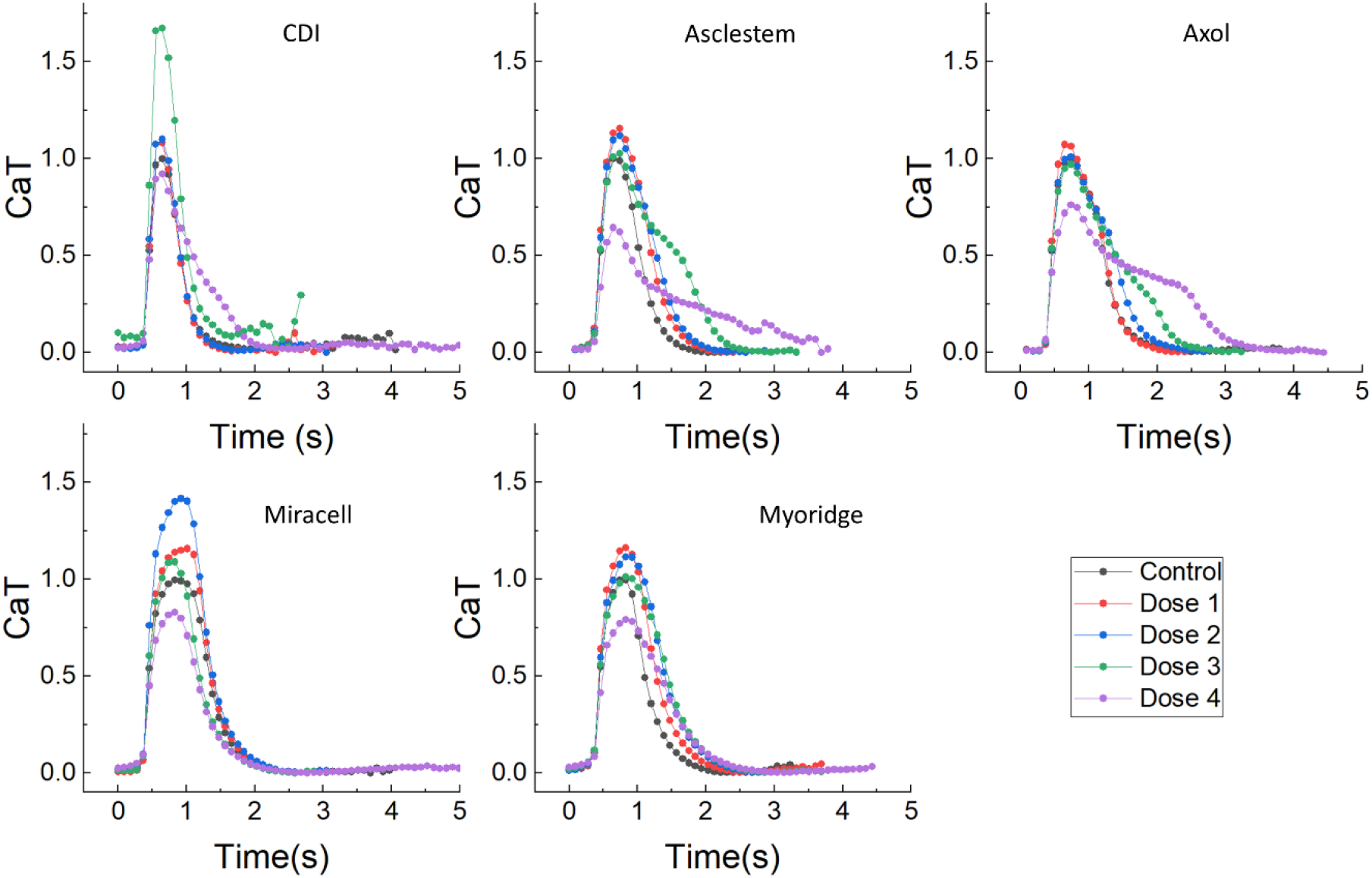
Effect of reduced calcium transient amplitude after treatment with Ranolazine.

**Figure S6.**
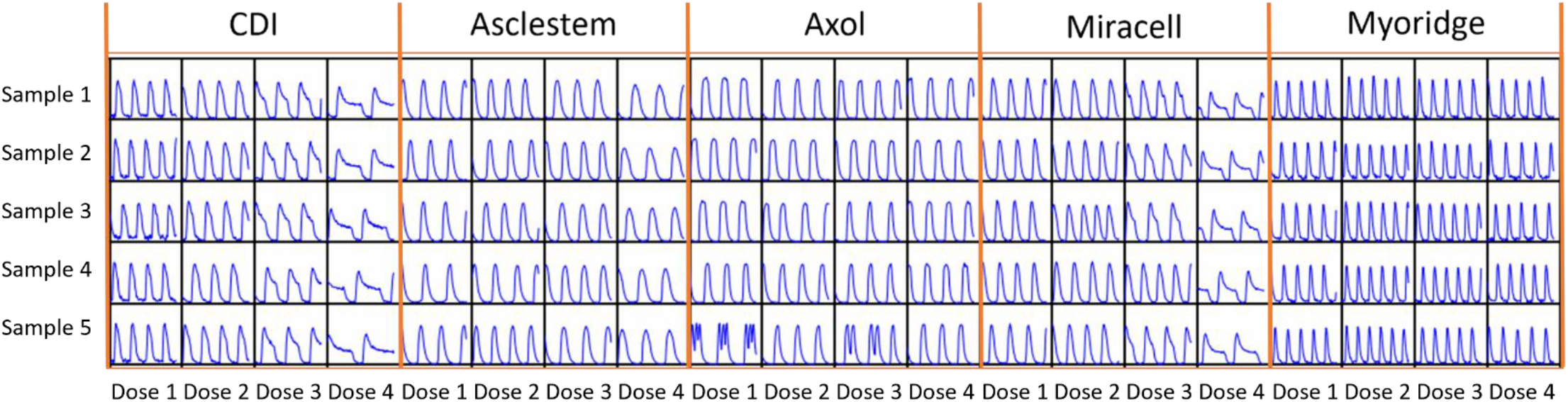
Quinidine-induced changes in raw calcium transient profiles (plots of calcium intensity vs. time). CCA analysis indicated reduced sensitivities of changing CaT profiles by increasing concentration of quinidine in Asclestem, Axol, and Myoridge. An overview of raw data of all samples with all doses clearly indicated the difference in quinidine sensitivity between CDI & Micrcell vs. Asclestem, Axpl, & Myoridge group.

**Figure S7.**
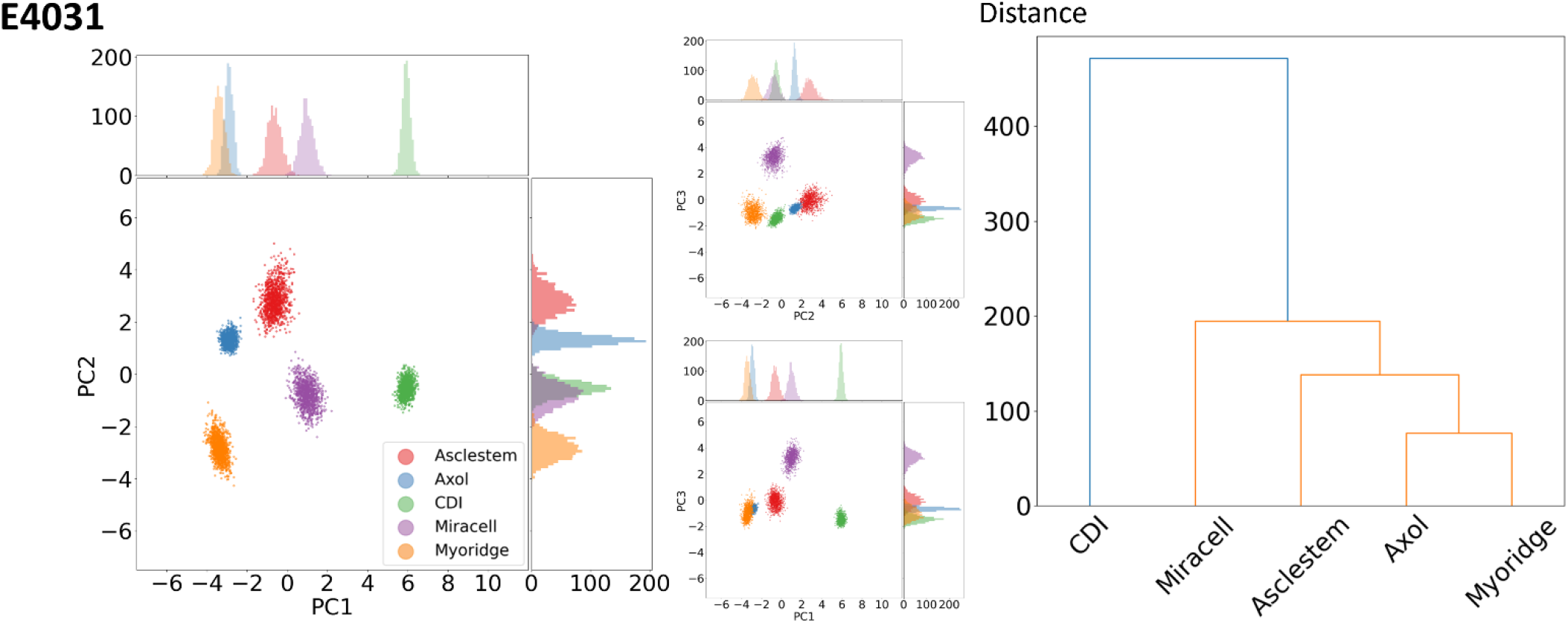
Cluster plot analysis and dendrogram of E4031 treated hiPSCM. E4031 is an experimental (research purpose only) class III antiarrhythmic drug that blocks hERG K+ channels. Cluster analysis of five cell lines treated with E4031 (upper left). Cluster analysis was generated by combining the six parameters and the four doses for each cell line into a matrix, resampling using Bayesian bootstrapping (1000) and performing dimensionality reduction by principal component analysis. The larger cluster analysis graph is a comparison of the first principal component (PC1) and second principal component (PC2). The two smaller cluster analysis graphs are a comparison of the second principal component (PC2) and third principal component (PC3) (top) and the first principal component (PC1) and third principal component (PC3) (bottom). Hierarchical cluster analysis (dendrogram, upper right) shows the hierarchical relationship between the five cell lines treated with E4031. Ward’s minimum variance method was used to calculate linkages. The colors represent different hierarchical clusters.

**Figure S8.**
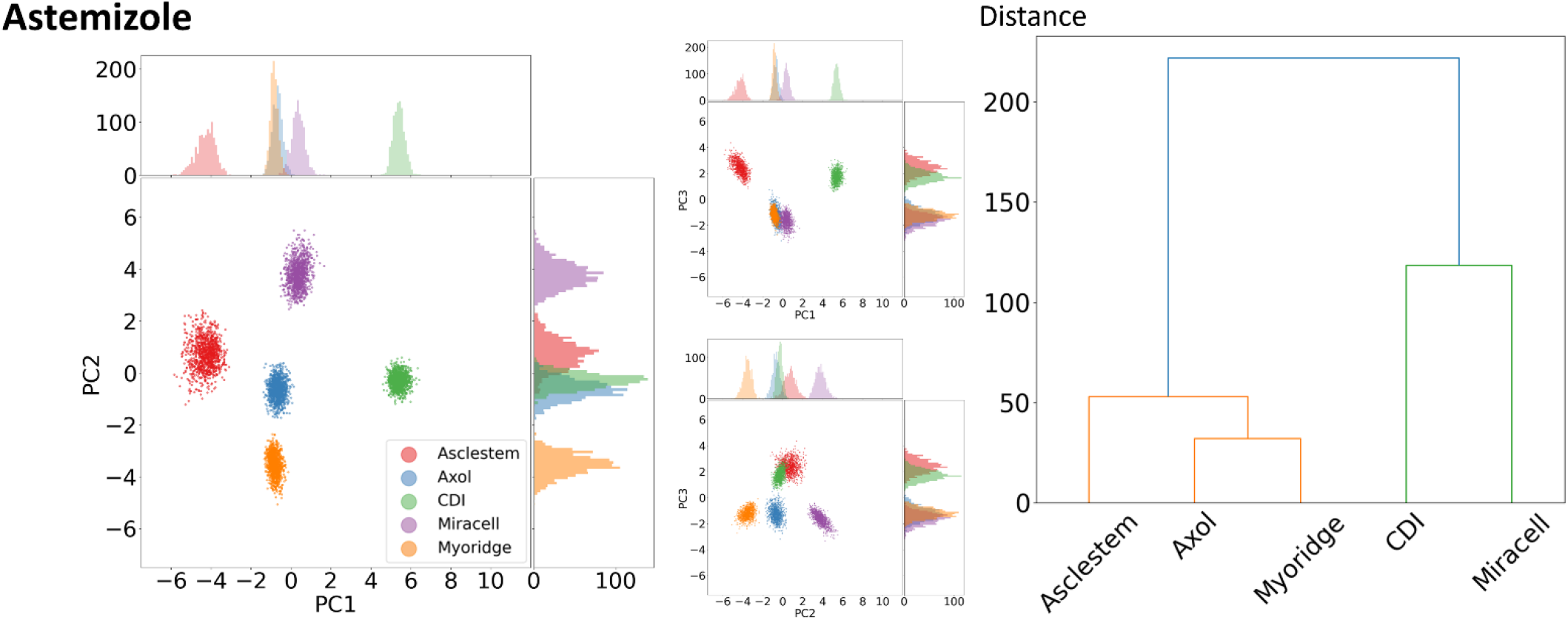
Cluster plot analysis and dendrogram of Astemizole treated hiPSCM. Astemizole is an antihistamine that blocks hERG K+ channels. The hierarchical cluster analysis divides the cells into two groups: 1) CDI and Miracell and 2) Asclestem, Axol and Myoridge.

**Figure S9.**
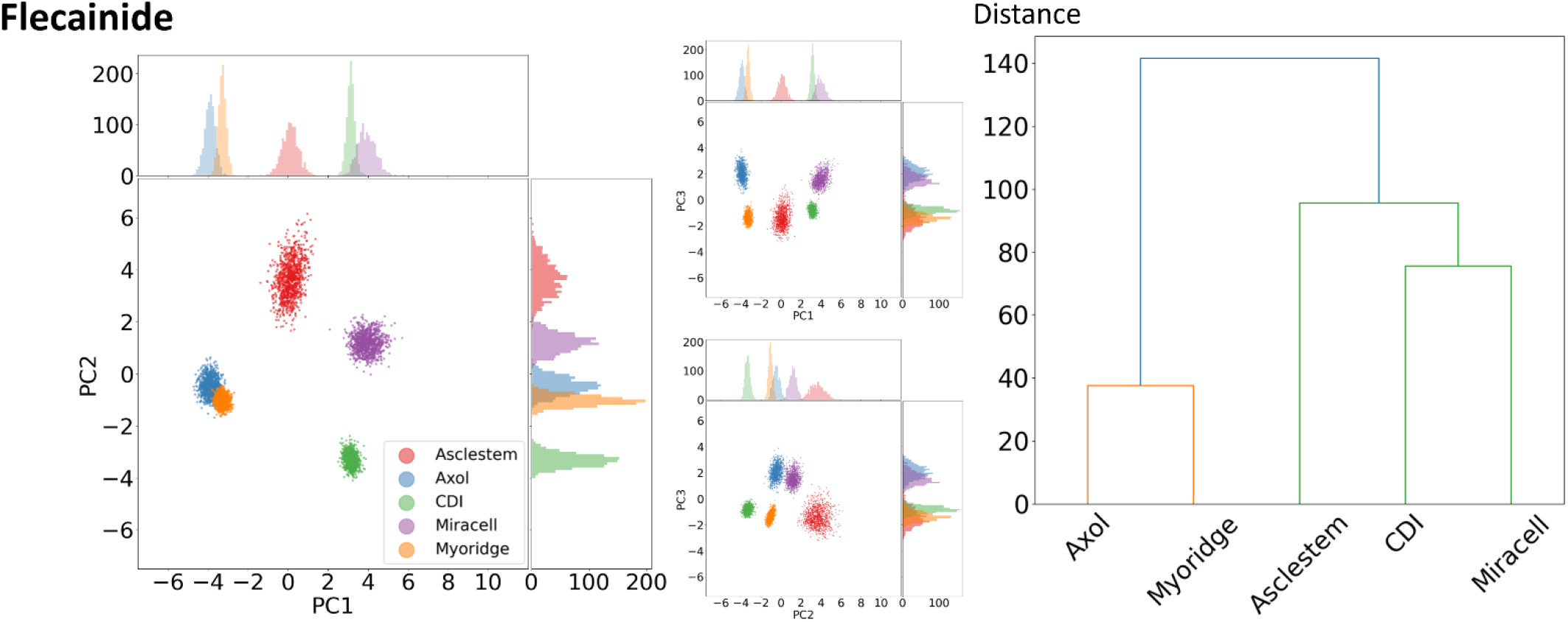
Cluster plot analysis and dendrogram of Flecainide treated hiPSCM. Flecainide is a class 1c antiarrhythmic drug that inhibits Na and K+ channels. Cluster analysis shows a close grouping of clusters for Axol and Myoridge. Hierarchical cluster analysis separates into two groups: 1) Axol and Myoridge, and 2) CDI, Miracell and Asclestem.

**Figure S10.**
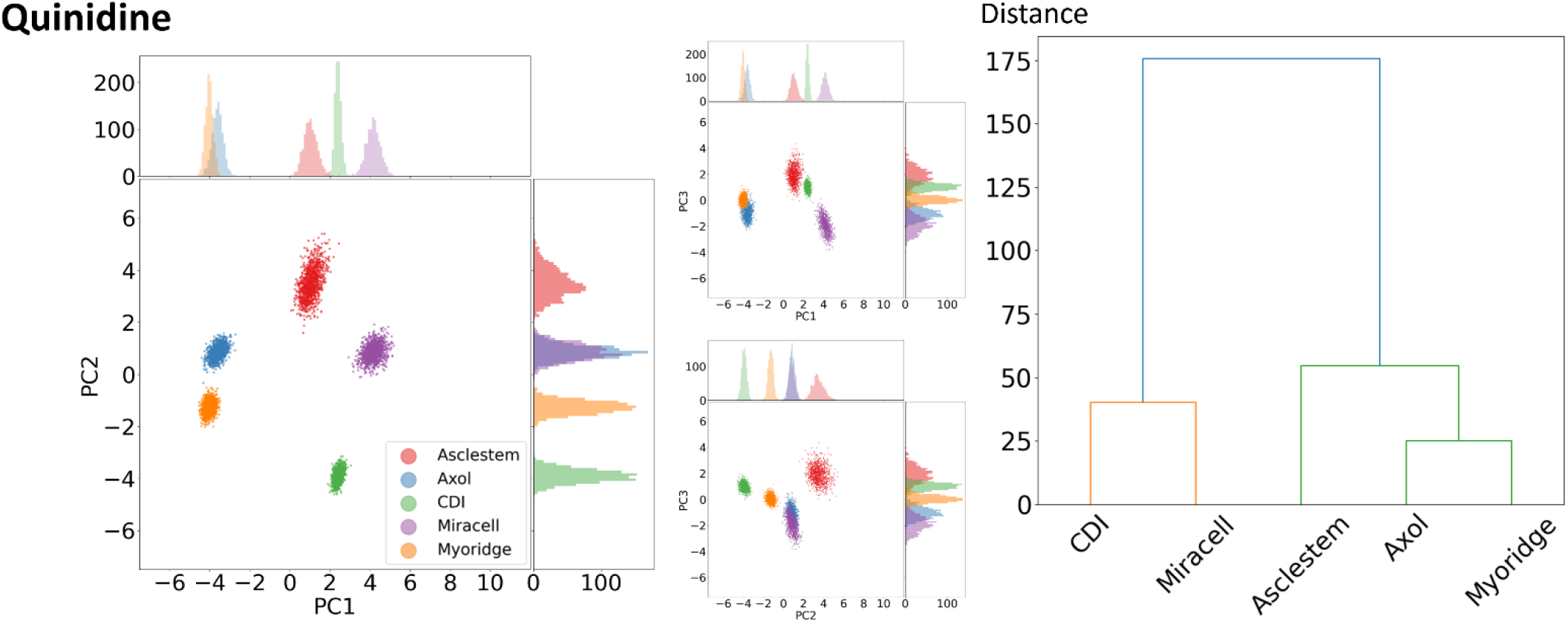
Cluster plot analysis and dendrogram of Quinidine treated hiPSCM. Quinidine is a class 1A antiarrythmic drug that blocks voltage-gated K+ channels. Cluster analysis shows a close grouping of clusters for Axol and Myoridge. Hierarchical cluster analysis separates into two groups, 1) CDI and Miracell, and 2) Asclestem, Axol and Myoridge.

**Figure S11.**
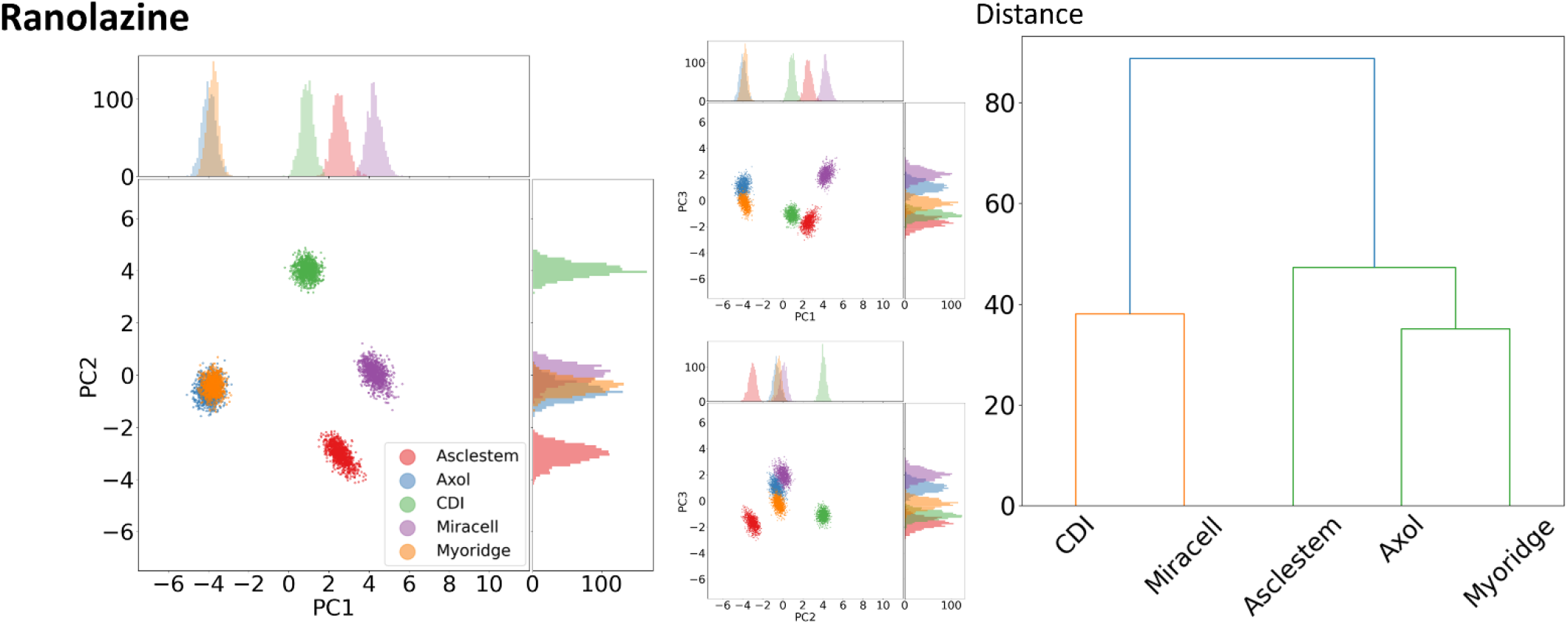
Cluster plot analysis and dendrogram of Ranolazine treated hiPSCM. Ranolazine inhibits late inward Na currents and K+ channels and is used to treat chronic angina. The radial plot (bottom left) shows ranolazine inhibits IKr and late INa. Cluster analysis shows similar locations of clusters for Axol and Myoridge. Hierarchical cluster analysis separates into two groups, 1) CDI and Miracell and 2) Asclestem, Axol and Myoridge.

**Figure S12.**
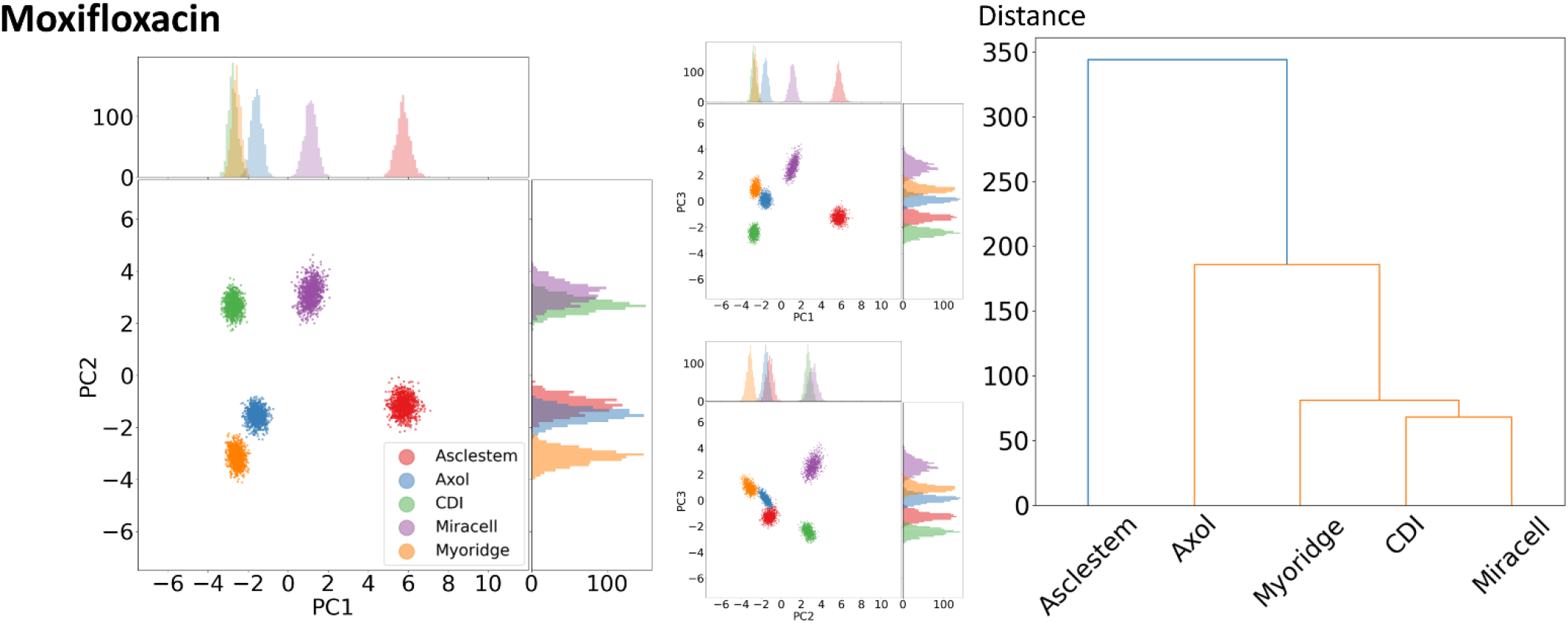
Cluster plot analysis and dendrogram of Moxifloxacin treated hiPSCM. Moxifloxacin is broad-spectrum antibiotic. Cluster analysis shows clusters for Axol and Myoridge are in proximity. The hierarchical cluster analysis separates Asclestem from CDI, Miracell, Myoridge, and Axol.

**Figure S13.**
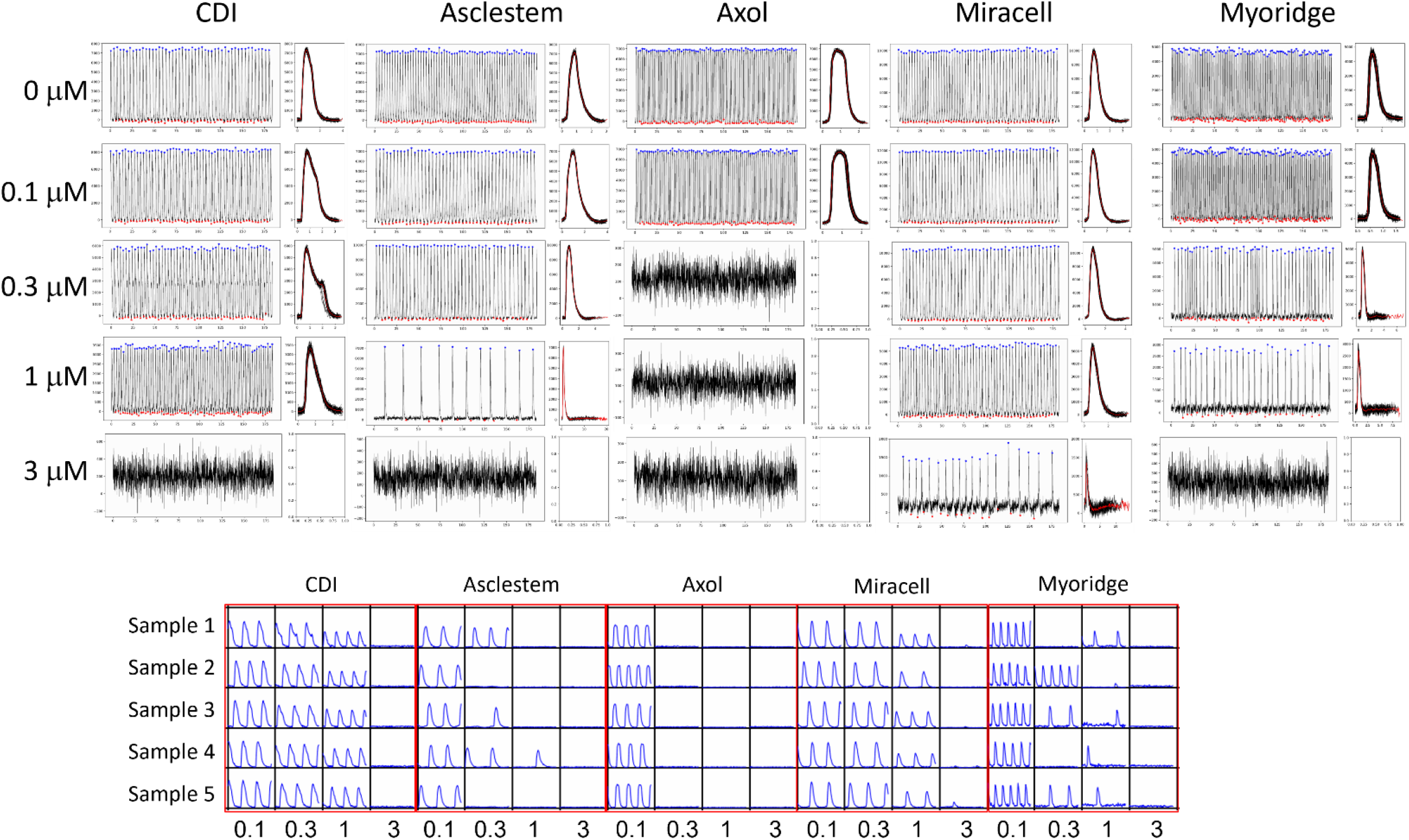
Effect of Terfenadine on different hiPSCM cell lines. Terfenadine slightly increased CDI’s CaT duration at 0.1-0.3 μM but not the other cell clines. Beat rates of Asclestem, Miracell, and Myoridge are reduced without changing their CaT profile significantly.

**Figure S14.**
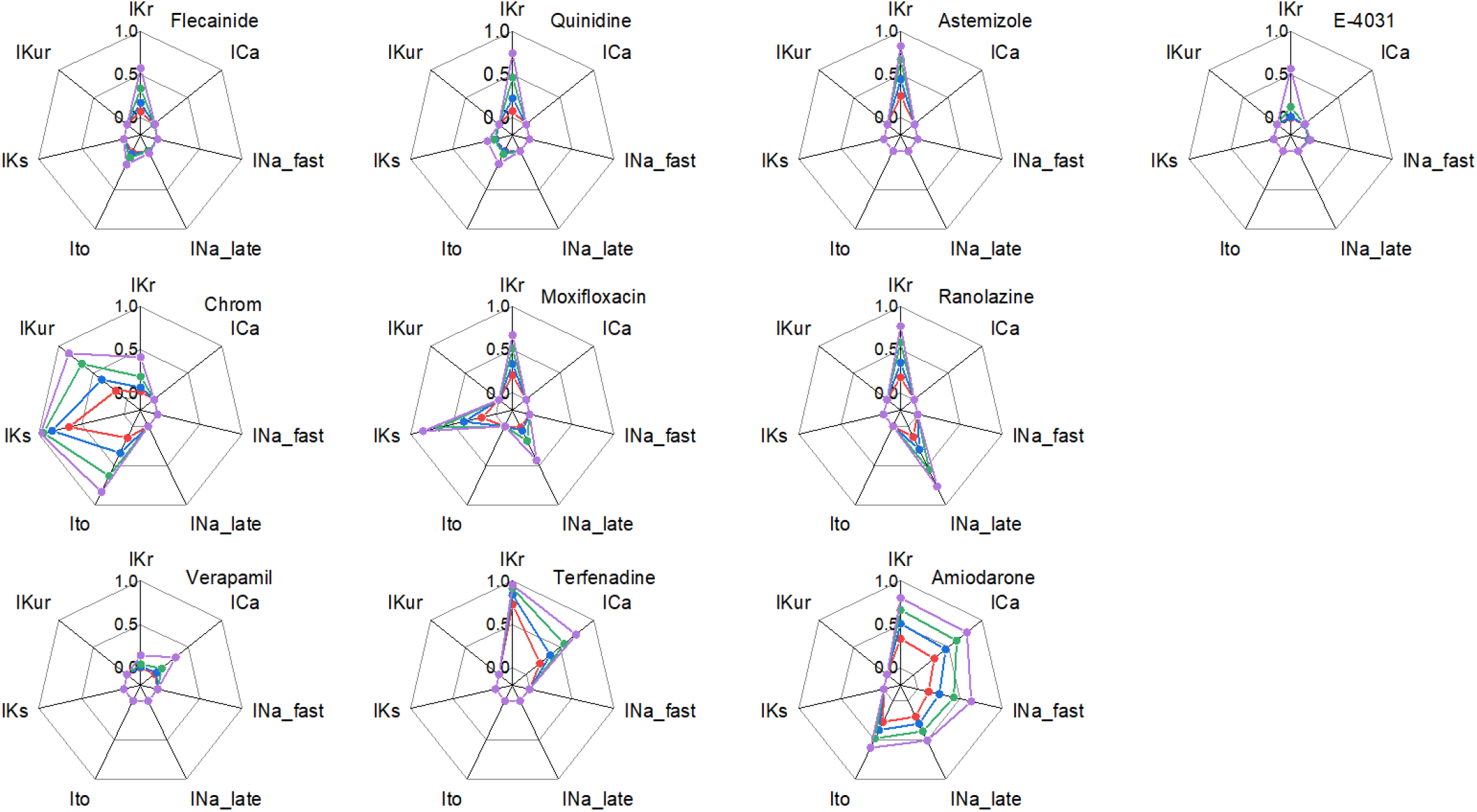
Expected degree of channel inhibitions. Inhibition ranges of ion channels are plotted 0.0 and 1.0 (0 to 100% inhibition). Red, blue, green, and purple represent dose 1, 2, 3, and 4, respectively. IKr: inward-rectifier potassium channel, ICa: L-type calcium channel, INa_fast: fast sodium channel, INa_late: late sodium channel, Ito: transient outward potassium channel, IKs: delayed rectifier potassium channel, IKur: ultra rapid potassium channel. Inhibition data were obtained from the FDA study [1].

## Footnotes

1. Blinova, K., et al., Comprehensive Translational Assessment of Human-Induced Pluripotent Stem Cell Derived Cardiomyocytes for Evaluating Drug-Induced Arrhythmias. Toxicol Sci, 2017. **155**(1): p. 234-247.

